# A flow cytometry-based screening platform for identifying candidate radiosensitizers targeting DNA repair

**DOI:** 10.64898/2026.08.25.747024

**Authors:** Christian Naucke, Gro Elise Rødland, Adrian Eek Mariampillai, Sissel Hauge, Linn Helena Steive, Ingvild Alida Bjerke, Lilian Lindbergsengen, Anne Serina Gilje Grøsvik, Vilde Siggerud, Karoline Kongsrud, Diana Savu, Trond Stokke, Randi G. Syljuåsen

**Author notes:** Correspondent author, Randi G. Syljuåsen, Department of Radiation Biology, Institute for Cancer Research, Norwegian Radium Hospital, Oslo University Hospital, Ullernchausseen 70, N-0379 Oslo, Norway.

## Abstract

Radiotherapy induces cytotoxic DNA damage, but activation of DNA repair pathways and cell-cycle checkpoints can limit therapeutic efficacy. Here, we developed a high-throughput, flow cytometry–based screening platform to identify compounds that inhibit radiation-induced DNA repair and checkpoint activation. Reh leukemia and A549 lung cancer cells were irradiated and screened against up to 700 bioactive compounds, with DNA damage persistence quantified by γH2AX levels across independent screens. Cell barcoding using Pacific Blue staining was incorporated to enable highly accurate quantification of γH2AX across treatment conditions. The platform yielded robust and reproducible results and supported multiparametric analysis, including assessment of G2 checkpoint activation by phospho-histone H3. Largely overlapping candidate radiosensitizers were identified in both cell lines, including the multi-kinase inhibitor 5-iodotubercidin and the PI3K/mTOR inhibitor omipalisib. Validation studies in lung cancer and glioblastoma models confirmed screen performance. Mechanistically, omipalisib reduced phosphorylation of the non-homologous end-joining protein DNA-PK, consistent with impaired double-strand break repair. Both compounds enhanced radiosensitivity in clonogenic survival assays. Notably, 5-iodotubercidin increased radiosensitivity in glioblastoma cells despite previous reports of radioprotective effects in normal brain tissue. Together, these findings establish a robust barcoded screening approach for identifying radiosensitizers that target DNA damage repair and checkpoint responses.

## Introduction

Radiotherapy is a cornerstone of cancer treatment, contributing to the cure of approximately 40% of patients(1). However, efficient DNA repair in tumor cells can lead to radioresistance and limit treatment effectiveness (2–4). Radiation-induced DNA damage is repaired through multiple pathways, including homologous recombination (HR) and non-homologous end-joining (NHEJ) for DNA double-strand breaks (DSBs), and base excision repair (BER) for single-strand breaks (SSBs) (5). In addition, activation of the G_2_ cell cycle checkpoint following irradiation provides additional time for repair prior to cell division, further enhancing cell survival (6).

To improve tumor radiosensitivity, combining radiotherapy with inhibitors of DNA repair or the G_2_ checkpoint has emerged as a promising strategy (7–9). In addition to established inhibitors targeting e.g. ATM, ATR, DNA-PK, or PARP (9), many emerging cancer drugs may also interfere with DNA repair or checkpoint control. Moreover, approved drugs developed for non-oncological indications could potentially be repurposed to enhance radiotherapy responses (10). In this context, compound libraries of novel and approved agents enable systematic screening to identify drugs that impair DNA repair after radiation.

Previous studies have employed microscopy-based high-throughput screening assays involving compound libraries and radiation to identify inhibitors of DNA repair (e.g. (11, 12)). While these approaches have provided valuable insights, they are often limited by time-intensive image acquisition and analysis, typically involving fewer cells and offering less accessible multiparameter analysis compared to flow cytometry-based methods. Recent advances now enable flow cytometry-based screening in 384-well formats, allowing rapid, multiparameter analysis of thousands of single cells per sample. Here, we present a novel flow cytometry screening platform designed to identify drugs that inhibit DNA repair and/or G_2_ checkpoint activation following irradiation. Using this method, we obtained robust results in two cancer cell lines and identified omipalisib and 5-iodotubercidin as inhibitors of radiation-induced DNA repair.

## Materials and methods

### Cell culture and irradiation

The B-lymphoid precursor cell line Reh was originally derived from a patient with acute lymphoblastic leukemia (13) and was kindly provided by Dr. M. F. Greaves (Imperial Cancer Research Fund Laboratories, London, UK) (14). Human A549 and H460 lung cancer cell lines and T98G glioblastoma cells were purchased from ATCC, while U251-MG glioblastoma cells were purchased from CLS Cell Lines Service GmbH (Germany). Reh cells were cultured in RPMI 1640 medium and the other cell lines in Dulbecco’s modified Eagle’s (DMEM) medium (both media from Life Technologies), at 37°C in a humidified atmosphere with 5% CO_2_. The media were supplemented with 10% fetal bovine serum (South America origin, Life Technologies) and 1% penicillin (10.000 U/ml)/streptomycin (10.000 μg/ml)(Life Technologies). All cell lines were authenticated by short tandem repeat profiling and regularly tested for *Mycoplasma* infection. Faxitron CP160 X-ray machine was used for irradiation (160 kV, dose rate: 1 Gy/min).

### Flow cytometry-based high-throughput screen

A detailed description of the flow cytometry-based high-throughput screening procedure is provided in Supplementary Methods. In brief, compound libraries (Enzo Target and Pathway, Selleckchem Cambridge Cancer, and BioMol Kinase Inhibitor libraries) were obtained through the Chemical Biology Platform and screened in 384-well plates. A549 lung cancer or Reh leukemia cells were seeded at 1 × 10⁵ cells per well and treated with compounds at a final concentration of 10 µM (1 µM for one Selleckchem library screen). Three parallel plates were processed per experiment: one non-irradiated control plate and two plates irradiated with 6 Gy. This dose induced robust γH2AX formation within 30 min with substantial resolution by 6 h, enabling detection of impaired DNA repair. Irradiated plates were harvested at 30 min and 6 h post-irradiation, and non-irradiated plates at 6 h. Cells were fixed in methanol and stored at −20°C until staining. Samples from the three plates were barcoded using differential Pacific Blue labeling and pooled into a single plate prior to antibody staining and analysis. Cells were stained with an anti-γH2AX antibody to quantify DNA damage. For assessment of G_2_ checkpoint abrogation in Reh cells, co-staining with anti-phospho-H3 was performed. DNA content was assessed using FxCycle Far Red staining. Data were acquired on a LSR II flow cytometer (BD Biosciences) equipped with a high-throughput sampler. Treatment groups were deconvoluted based on Pacific Blue intensity. Median γH2AX levels and the percentage of phospho-H3-positive mitotic cells were quantified. Candidate compounds affecting DNA repair were identified by calculating the ratio of γH2AX levels at 6 h versus 30 min after irradiation, with subtraction of non-irradiated controls (γH2AX_ratio_= (γH2AX_6Gy,6h_ – γH2AX_0Gy,6h_)/ (γH2AX_6Gy,30min_ – γH2AX_0Gy,6h_)). Compounds abrogating the G_2_ checkpoint were identified based on increased mitotic entry at 6h after irradiation without affecting non-irradiated samples. Thresholds for hit selection were defined empirically based on population distributions.

### Flow cytometry analysis of γH2AX and phospho-53BP1

Flow cytometry analysis of individual samples (Figures 5 and 6) was performed as described previously (15). Briefly, cells were fixed in 70% ethanol and stained with antibodies against γH2AX (Millipore, clone JBW301, 1:500) and phospho-53BP1 (Cell Signaling Technology, #2675S, 1:100), followed by Alexa Fluor™ 488- and 568-conjugated anti-mouse and anti-rabbit secondary antibodies (Invitrogen, 1:1000), in flow buffer (0.1% Igepal CA-630, 6.5 mM Na₂HPO₄, 1.5 mM KH₂PO₄, 2.7 mM KCl, 137 mM NaCl, 0.5 mM EDTA, pH 7.5) containing 4% non-fat milk powder. Barcoded cells from a separate sample, stained with Alexa Fluor™ 647 succinimidyl ester (Invitrogen), were added to each sample prior to antibody staining and served as an internal standard; populations were distinguished by gating during analysis. Samples were analyzed using an LSRII flow cytometer (BD Biosciences) and processed with FACSDiva and FlowJo software (BD Biosciences).

### Western blotting

Western blotting was performed as described previously (15). Cells were lysed in ice-cold TX-100 buffer (100 mM NaCl, 50 mM Tris pH 7.5, 2 mM MgCl₂, 0.5% Triton X-100) supplemented with Benzonase (100 U/ml), EDTA-free protease inhibitors, and phosphatase inhibitors. After incubation at 4°C, reducing sample buffer was added and samples were boiled (95°C, 5 min). Proteins were separated using Criterion Stain-Free TGX gels and transferred to nitrocellulose membranes. Antibodies are listed in Supplementary Table 1. Chemiluminescent signals were acquired on a ChemiDoc MP system and quantified using Image Lab software. Membranes were stripped using ReBlot Plus Mild Stripping Solution for reprobing.

### Clonogenic survival assay

Cells were seeded at low density (250-4000 cells per 6-cm dish), in triplicate (experiments with omipalisib) or sextuplicate (experiments with 5-iodotubercidin), and treated 20-24 h later with drugs and radiation. Omipalisib and 5-iodotubercidin were added 30 min prior to irradiation. Colonies were allowed to grow for 12-14 days, followed by fixation in 70% ethanol and staining with methylene blue. Colonies containing >50 cells were counted. The survival fraction was calculated as: (number of colonies formed / number of cells seeded) for treated samples, normalized to the corresponding value for untreated controls.

## Results

### Design of the screening method

To identify compounds that inhibit DNA repair after radiation, we designed a flow cytometry-based screening method (Figure 1A-C). DNA repair is assessed by an antibody to the DNA damage marker γH2AX. At 30 minutes the γH2AX level is high due to the radiation-induced DSBs, but at 6 hours it has decreased due to DNA repair. For drugs that inhibit DNA repair, we expect higher levels of γH2AX remaining at 6 hours (Figure 1C, upper panels). Furthermore, by inclusion of an antibody to the mitotic marker phospho-H3 S10, the screening method can simultaneously identify compounds that abrogate the radiation-induced G_2_ checkpoint (Figure 1B-C, bottom panels). At 6 hours after radiation there are no mitotic cells due to activation of the G_2_ checkpoint, but mitotic cells will be present when a drug abrogates the G_2_ checkpoint (Figure 1C, bottom panels). The method can thus identify compounds that directly inhibit radiation-induced DNA repair pathways, and/or indirectly inhibit DNA repair by abrogating the G_2_ checkpoint. A key feature of our method is the use of Pacific Blue-based barcoding, which enables accurate detection of subtle differences in DNA repair while reducing antibody consumption. Each of the three plates (0 Gy, 6 Gy 0.5 h, 6 Gy 6 h) is labeled with distinct Pacific Blue concentrations and then pooled into a single plate for antibody staining, thus eliminating variation from antibody staining. During the final analysis, samples are separated based on their Pacific Blue signal (Figure 1B).

**Figure 1.**
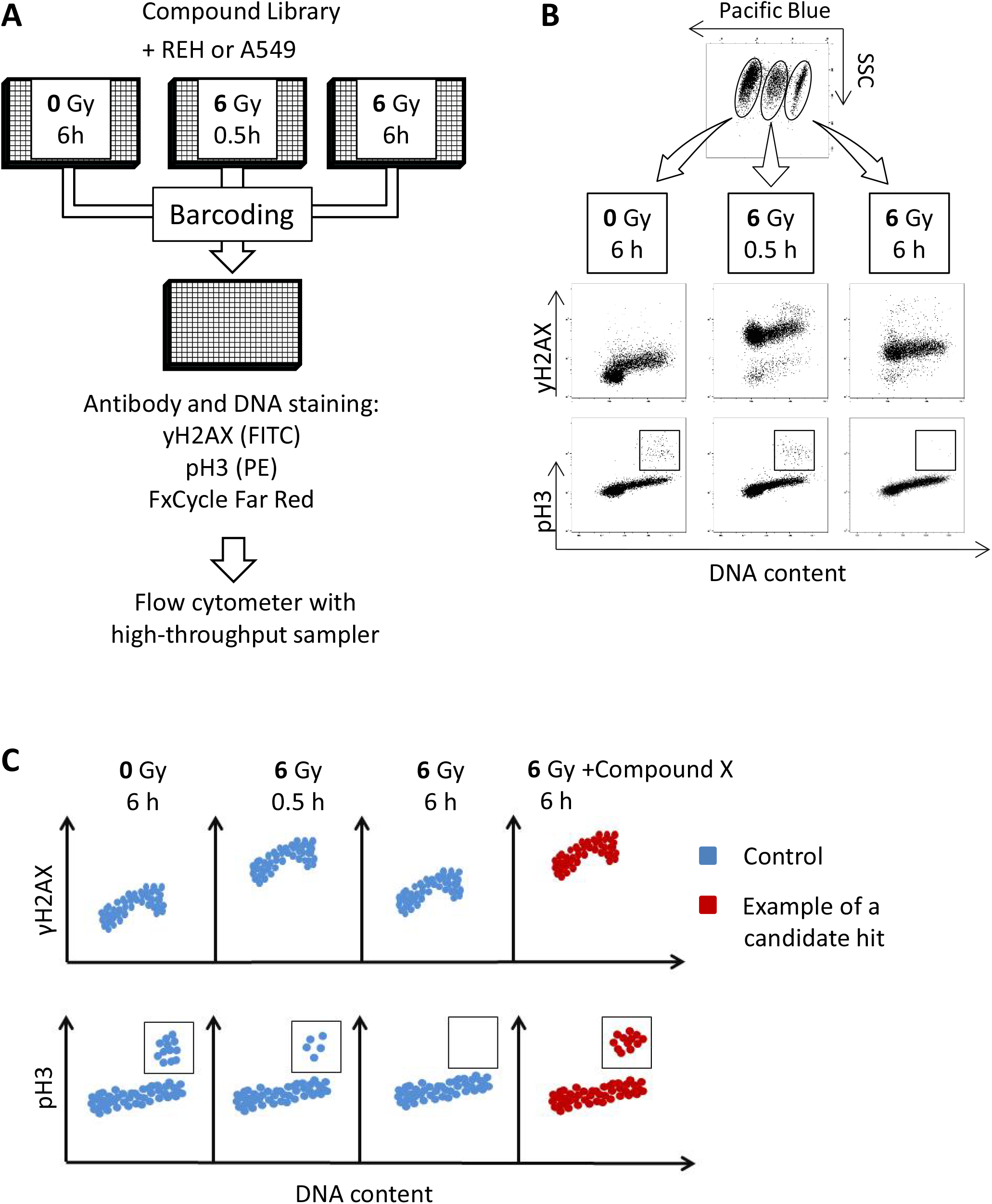
Screen layout. **A.** Cells were seeded in three 384-well plates with compounds. Two plates were irradiated (6 Gy) and harvested at 30 min and 6 h; the third (0 Gy) was harvested at 6 h. Samples were barcoded, pooled, stained for γH2AX (FITC), phospho-H3 S10 (PE), and DNA (Far Red), then analyzed by high-throughput flow cytometry. **B.** Barcoded samples were demultiplexed and analyzed to generate scatter plots of γH2AX versus DNA content and phospho-H3 versus DNA content, with mitotic cells appearing in distinct regions. **C.** Example plots illustrating expected results after irradiation alone (blue) and with a candidate hit (compound X; red).

### Screening of 357 compounds in Reh leukemia cells

Reh leukemia cells were used to optimize the screen because they grow in suspension, allowing high yields of single cells in 384-well plates. They have also been successfully employed in our previous flow cytometry screens for replication stress regulators (15), and they show robust induction of γH2AX and G_2_ checkpoint activation after irradiation (16). We performed two independent screens in Reh cells using the γH2AX endpoint with 357 compounds from the Enzo Target & Pathway and Biomol Kinome libraries. SAHA, a histone deacetylase (HDAC) inhibitor previously reported to inhibit DNA repair after irradiation (17), scored as a candidate hit (Figure 2A), confirming the validity of the screening method. To identify additional hits, we calculated the γH2AX ratio at 6 hours relative to 30 minutes post-irradiation for all compounds, as described in Materials and methods. Most compounds did not significantly alter the γH2AX ratio compared to irradiated controls, indicating no effect on DNA repair (Figure 2B, main peak). However, a subset of compounds showed higher ratios, consistent with impaired repair (Figure 2B, right of stippled line). Compounds scoring in both screens were classified as candidate hits (Figure 2C, quadrant Q1), including eleven HDAC inhibitors, two protein phosphatase (PP1/PP2A) inhibitors, and the adenosine kinase and multi-kinase inhibitor 5-iodotubercidin (Figure 2D and Table S2). Of note, a high γH2AX ratio could theoretically arise not only from increased γH2AX levels at 6 hours after irradiation, but also from reduced induction at 30 minutes. However, most of the identified candidate hits clearly showed elevated γH2AX levels at 6 hours, as evident when comparing the raw data for controls, the bulk of non-hit compounds, and the identified hits (Figure 2E).

**Figure 2.**
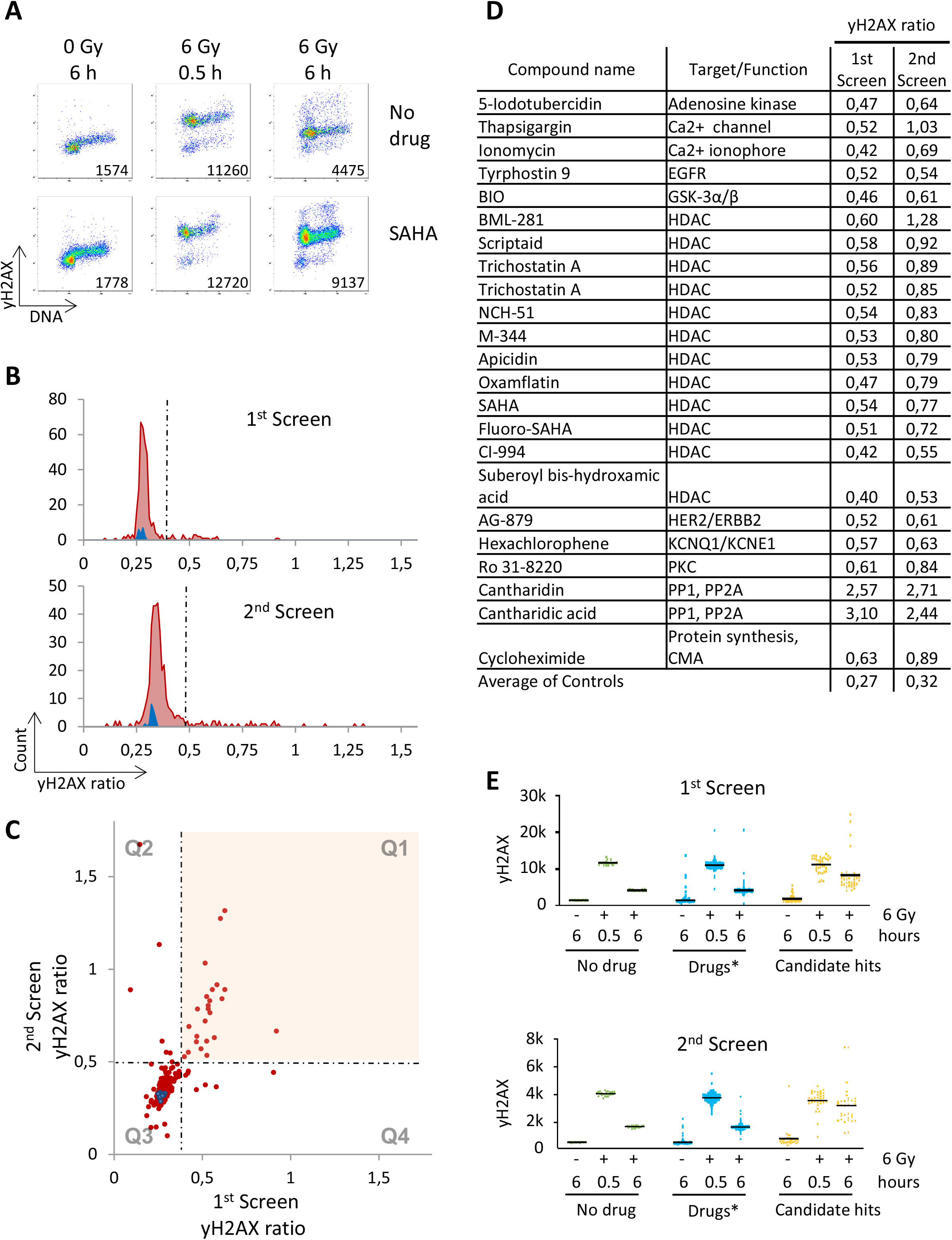
Screening results for 357 compounds from the Enzo pathway and Biomol kinome libraries in Reh cells (γH2AX endpoint). **A.** Example of scatter plots of γH2AX versus DNA content (FxCycle Far Red) from single wells, as in Figure 1A, B. Controls treated with radiation only are shown in the top panel, and cells treated with the HDAC inhibitor SAHA in the bottom panel. Numbers indicate median γH2AX levels. **B.** Histograms of the γH2AX ratio, calculated as (γH2AX_6Gy,6h_-γH2AX_0Gy,6h_)/ (γH2AX_6Gy,30min_-γH2AX_0Gy,6h_), for each compound (red) and for radiation-only controls (blue). Compounds with ratios above the dashed line were considered candidate hits. Top: first screen; bottom: second screen. **C.** Comparison of γH2AX ratios between the two screens. Each compound and control sample is plotted, with results divided into four quadrants (Q1–Q4). Q1 contains hits scoring in both screens; Q2 and Q4 contain hits scoring in one screen only; Q3 contains controls and non-hits. **D.** List of compounds scoring as hits in both screens (quadrant Q1 in panel C). **E.** Raw ãH2AX signals for candidate hits (yellow) compared with all other compounds (Drugs*) and controls (No drug). One datapoint in the first screen exceeding 30,000 was excluded from this plot for visualization purposes.

In the second screen, we also included phospho-H3 staining to assess G_2_ checkpoint abrogation. In untreated samples, ∼2% of cells were mitotic, whereas irradiation reduced this to <0.05% at 6 hours (Figure S1A–B). A few compounds caused a high mitotic fraction (3–12%) without irradiation, indicating mitotic arrest; these included known regulators such as nocodazole (Figure S1B, top panel, and Table S2). Candidate hits for G_2_ checkpoint abrogation were defined as compounds that did not increase mitotic index in non-irradiated samples (∼2%) but showed >1% mitotic cells at 6 hours after 6 Gy (Figure S1C). Four compounds met these criteria: MG-132, sanguinarine chloride, pimozide, and rapamycin (Figure S1D).

### Applying the screening method to A549 lung cancer cells

Radiotherapy is used more frequently to treat solid tumors than leukemia. We chose to test our screening method in non-small cell lung cancer (NSCLC), as radiotherapy is commonly used for advanced NSCLC and DNA repair inhibitors could potentially provide substantial benefit. We applied the screen to A549 NSCLC cells, which are relatively radioresistant, likely due to *RAS* mutation (18). Although typically adherent, A549 cells can grow in suspension (19), enabling adaptation to the screening format. Two screens were performed with the same 357-compound libraries as above using the γH2AX endpoint. (Phospho-H3–based assessment of mitotic entry was not performed in A549 cells because the mitotic index declined during the initial six hours after seeding in suspension.) SAHA scored as a candidate hit in the A549 screens (Figure 3A), consistent with the results in Reh cells (Figure 2A). Calculation of γH2AX ratios revealed that most compounds did not affect DNA repair, but a subset showed increased ratios, indicating inhibition. Compounds scoring in both screens were classified as candidate hits (Figure 3B), and showed substantial overlap with screens in Reh cells (Figure 3C; Figure S2). Shared hits included 10 of 11 HDAC inhibitors, two PP1/PP2A inhibitors, anisomycin, and 5-iodotubercidin. Some hits were cell-line specific; for example, sorafenib tosylate (a known radiosensitizer) scored in A549 but not Reh, whereas thapsigargin scored in Reh only.

**Figure 3.**
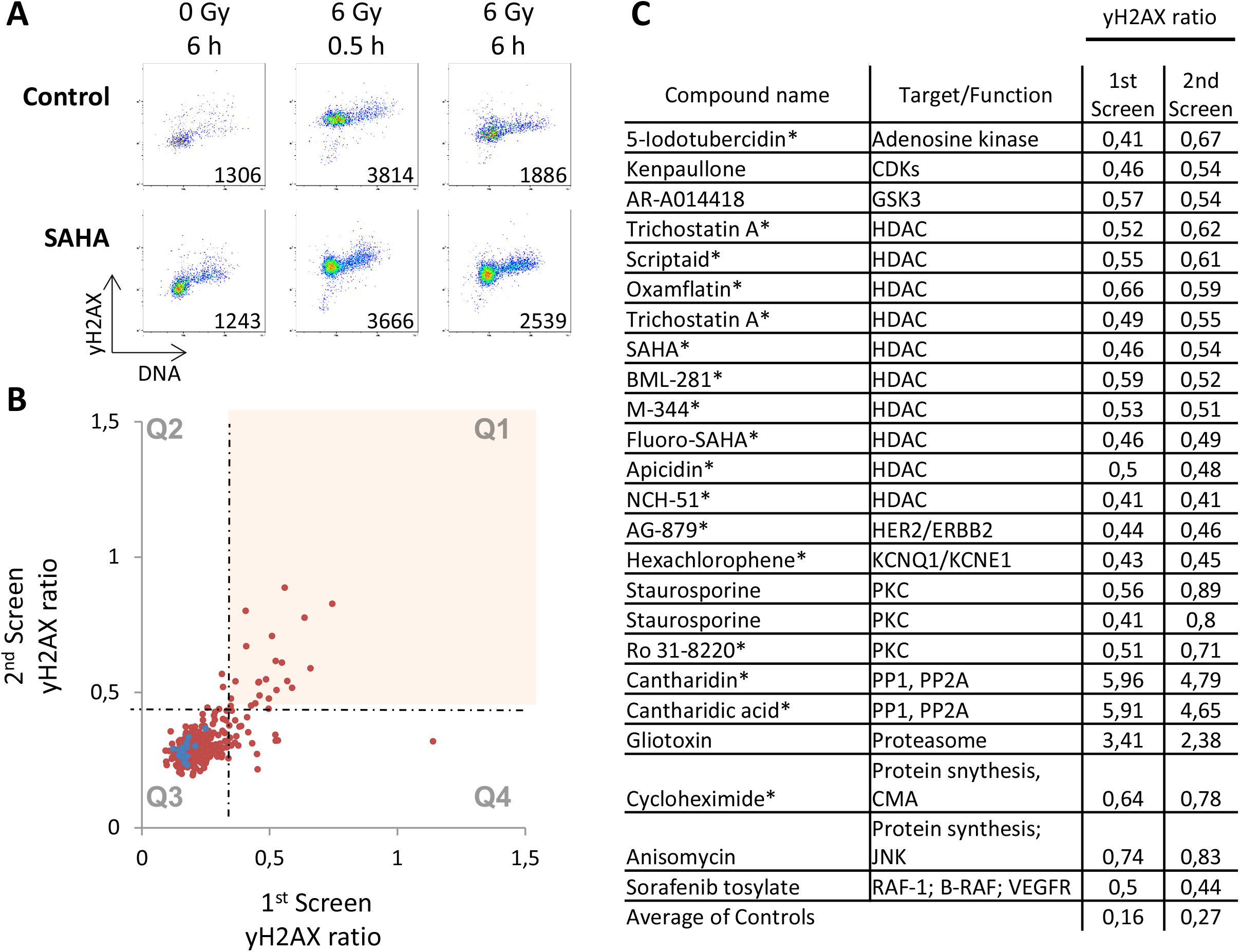
Screening results for 357 compounds from the Enzo pathway and Biomol kinome libraries in A549 cells. **A.** Example of scatter plots, as in Figure 2A, in A549 lung cancer. **B.** Comparison of γH2AX ratios between two screens, as in Figure 2C, in A549 cells. **C.** List of compounds scoring as candidate hits in both screens (compounds in quadrant Q1 in panel B). Compounds labeled * also scored as candidate hits in Reh cells (Figure 2D).

We next screened A549 cells with 384 cancer-related compounds from the Selleck library, performing two screens, at 10 µM and 1 µM (Figure 4A). The 1 µM screen was included because many compounds are likely cytotoxic at 10 µM and may kill cancer cells even without radiation, particularly when exposure times exceed the 6-hour duration of the short-term assay. Candidate hits scoring in both screens included 10 HDAC inhibitors, 3 CDK inhibitors, 2 mTOR inhibitors, (among them the dual PI3K/mTOR inhibitor omipalisib, GSK2126458), and 2 inhibitors of nucleotide metabolism (Figure 4B). For comparison, the library contained 13 HDAC inhibitors, 7 CDK inhibitors, 13 mTOR inhibitors and 12 DNA/RNA synthesis inhibitors. These results reinforce HDAC inhibitors as major hits, although negative results should be interpreted cautiously as some library compounds may lack activity. Notably, the two nucleotide metabolism inhibitors, clofarabine and cladribine, are previously known to inhibit DNA repair (20), further confirming the validity of our screen.

**Figure 4.**
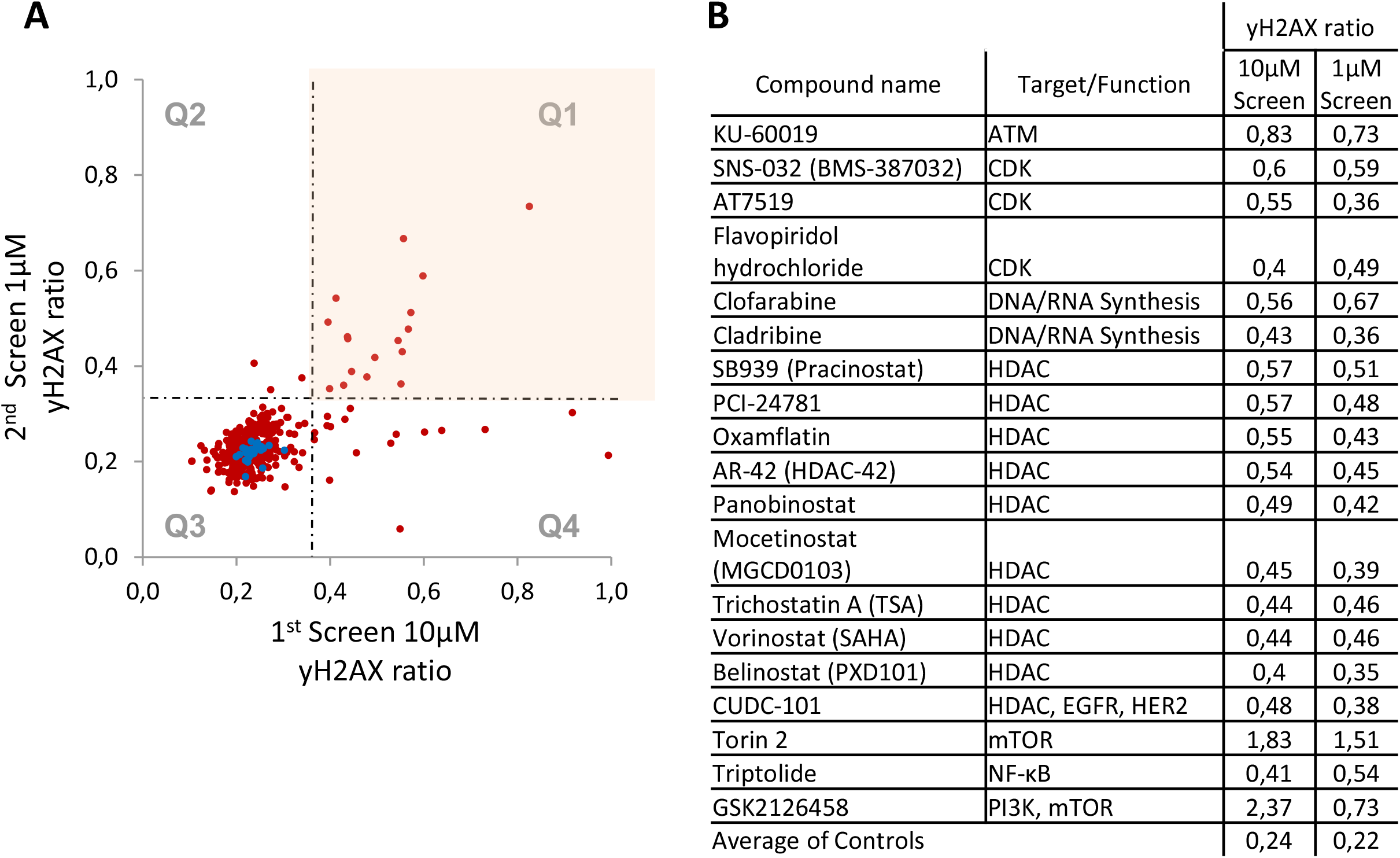
Screening results for 384 compounds from the Cancer Selleck Cambridge library in A549 cells. **A.** Comparison of ãH2AX ratios between two screens, similarly as in Figure 2C, in A549 cells with the Cancer Selleck Cambridge library. Compound concentrations were 10 μM (first screen) and 1 μM (second screen). **B.** List of compounds scoring as candidate hits in both screens (compounds in quadrant Q1 in A).

As mentioned above, a higher γH2AX ratio may reflect either increased γH2AX levels at 6 hours post-irradiation, consistent with impaired DNA repair, or reduced γH2AX levels at 30 min, indicating deficient early induction. To assess the latter, we examined γH2AX levels at 30 min after 6 Gy irradiation in the Selleckchem library screens. Most compounds did not affect early γH2AX induction (Figure S3A); however, a few compounds caused markedly reduced induction (Figure S3A,B). Based on this endpoint, ATM and DNA-PK inhibitors emerged as top hits inhibiting γH2AX induction (Figure S3C), consistent with their established roles in H2AX phosphorylation. Additional hits included mTOR and other PI3K inhibitors, including omipalisib (Figure S3C).

### Omipalisib induces radiosensitization and inhibits radiation-induced phosphorylation of DNA-PK

To validate and explore screen results, we repeated the screen assay with compounds acquired from an independent source than the compound libraries. We first focused on omipalisib, which has shown tolerability in Phase 1 clinical testing (21) and may play a role in radiosensitization and DNA repair (22, 23), although these effects have not been extensively studied. Consistent with the screen results and defective DNA repair, flow cytometry analysis of γH2AX showed that 0.5 μM omipalisib caused elevated γH2AX levels at 6 hours, while slightly reducing the levels at 30 minutes, after irradiation in A549 cells as well as in another lung cancer cell line, H460 (Figure 5A-C and S4A). Omipalisib also enhanced radiosensitivity in both lung cancer cell lines, as measured by clonogenic survival assays (Figure 5D). Immunoblotting of DNA damage signaling events revealed that omipalisib strongly suppressed radiation-induced phosphorylation of DNA-PK at S2056 (Figure 5E-F), suggesting inhibition of NHEJ repair. These results support and further extend the results of a previous study that showed radiosensitization of HeLa cells by omipalisib, and that omipalisib inhibits DNA-PKcs phosphorylation and activity (22). We conclude that omipalisib is a potent inhibitor of DNA-PK and this likely mediates its radiosensitizing properties, in agreement with the previous study (22).

**Figure 5.**
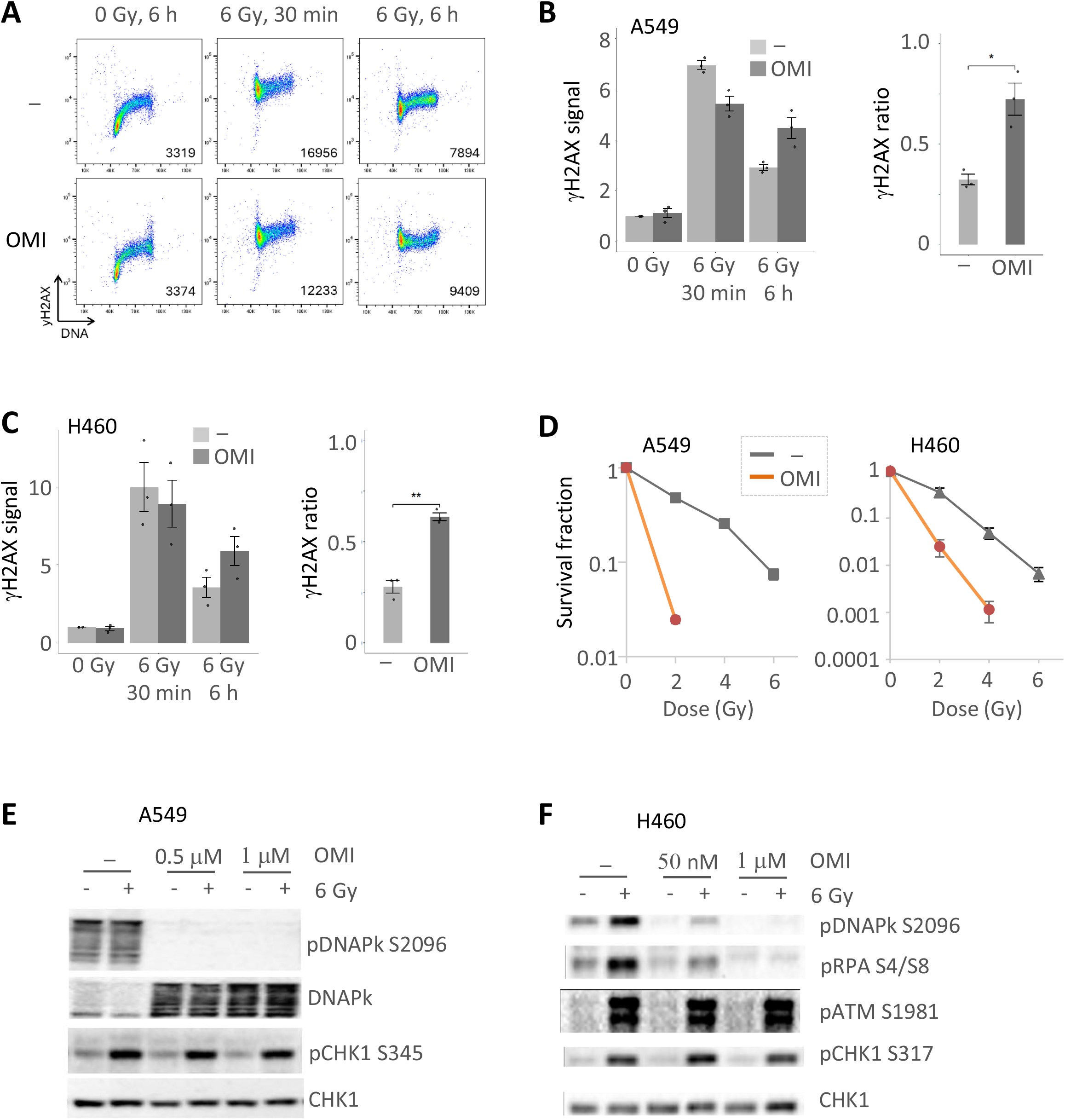
Omipalisib suppresses phospho-DNAPK and increases radiosensitivity in lung cancer cells. **A.** Scatter plots of γH2AX versus DNA content in A549 cells treated with radiation (6 Gy) and 0.5 μM omipalisib (OMI). Numbers indicate γH2AX median values. **B.** Quantification of γH2AX median values from experiments as in A. Values are shown relative to the non-treated sample. The γH2AX_ratio_ was calculated as (γH2AX_6Gy,6h_ – γH2AX_0Gy,6h_)/ (γH2AX_6Gy,30min_ – γH2AX_0Gy,30min_) n=3. Error bars represent the standard error of the mean (SEM), and each dot represents an individual experiment. *p<0.05, **p<0.01 (two tailed Student t test). **C.** Similar as in B for H460 cells. **D.** Clonogenic survival of A549 (left) and H460 (right) cells treated with omipalisib (0.5μM) and the indicated doses of radiation. Error bars: SEM (n=3). **E.** Immunoblots of A549 cells treated with omipalisib and radiation, as indicated. Cells were harvested 1 hour after treatment. The two upper blots (pDNAPk and DNAPk) were run on a separate part of the gel using the same samples as the two lower blots (pCHK1 and CHK1). **F.** Immunoblots of H460 cells treated with omipalisib and radiation, as indicated. Cells were harvested 1 hour after treatment. The pDNAPk, pRPA, and pATM blots were run on the same gel, while the pCHK1 and CHK1 blots were run on a separate gel in parallel using the same samples. Uncropped blots are shown in Supplementary Figure S5A (A549) and S5B (H460).

### Inhibition of DNA repair and radiosensitization by the multi-kinase inhibitor 5-iodotubercidin

We next explored the candidate hit 5-iodotubercidin, a multi-kinase inhibitor that potently targets Haspin, adenosine kinase and the dual-specificity tyrosine-regulated kinase (DYRK) and Cdc2-like (CLK) kinase families (∼10–100 nM), and more weakly inhibits Casein kinase 1 and additional kinases such as phosphorylase kinase and protein kinase A (PKA) (∼0.5–5 µM) (24–26). Flow cytometry analysis of γH2AX following treatment of A549 with radiation and 10 µM 5-iodotubercidin confirmed the screen results (Figure 6A, B and S4B), suggesting that this drug suppresses DNA repair in lung cancer cells. Since 5-iodotubercidin was previously shown to be radioprotective in normal mouse and rat brain tissue (27, 28), we wanted to investigate its potential radiosensitizing effects on brain tumor cells. To this end, we studied the effects of 5-iodotubercidin and radiation in two glioblastoma cell lines, U251 and T98G. Flow cytometry analysis showed that 0.5 μM 5-iodotubercidin led to elevated γH2AX levels at 24 hours after irradiation in both cell lines, suggesting defective DNA repair (Figure 6C, D). Consistently, levels of another marker for DSBs, p-53BP1, were also elevated at 24 hours (Figure S4C). Furthermore, immunofluorescence microscopy showed foci of γH2AX and p-53BP1 at 24 hours, in line with unrepaired DNA double strand breaks (Figure S4D). 5-iodotubercidin also reduced clonogenic survival after irradiation (Figure 6E), indicating radiosensitizing effects on glioblastoma cell lines. Immunoblot analysis of DNA damage signaling revealed that 5-iodotubercidin partly suppressed radiation-induced phosphorylation of CHK1 at S345 (Figure 6F), which may contribute to its radiosensitizing effect.

**Figure 6.**
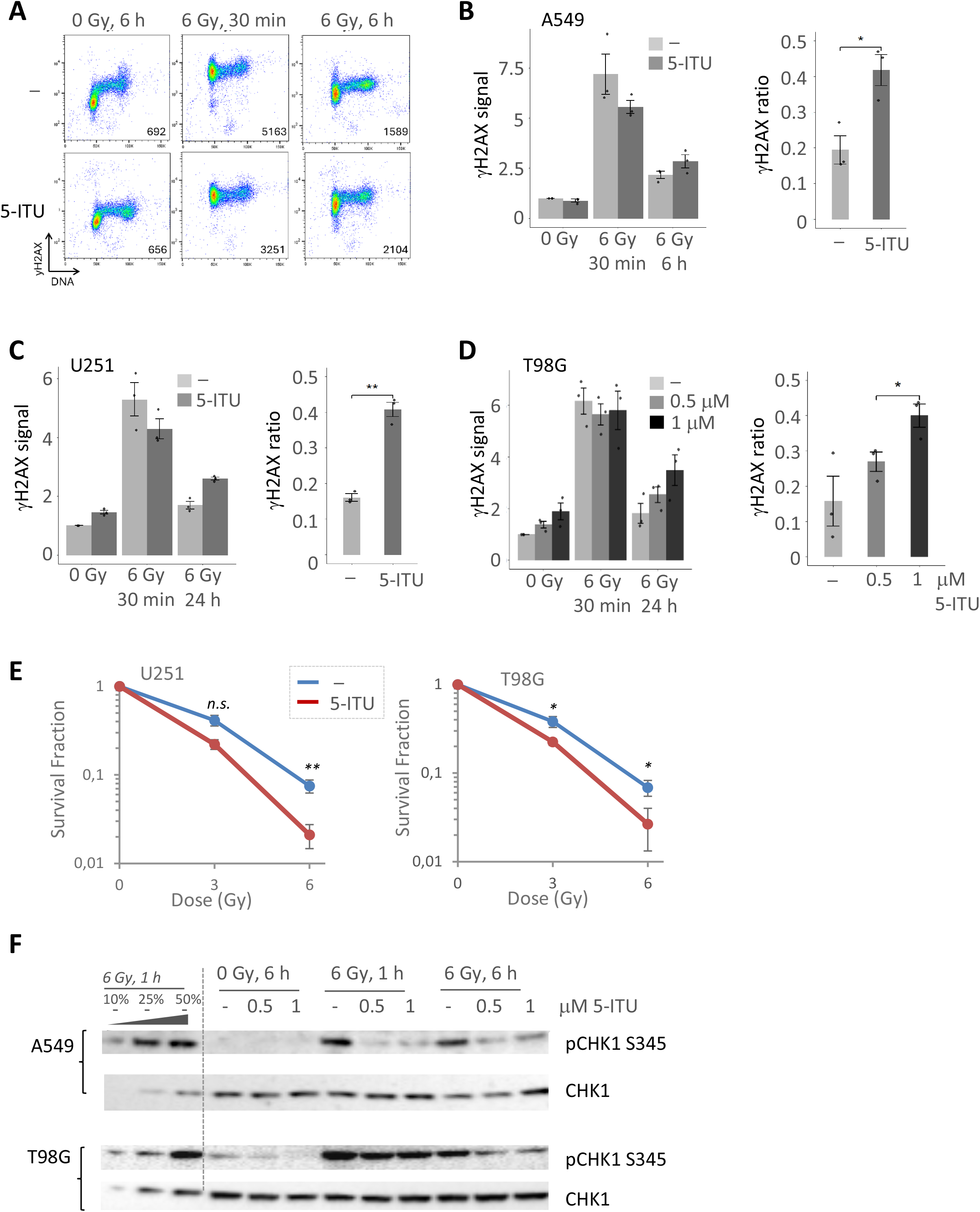
5-iodotubercidin suppresses DNA repair and increases radiosensitivity in glioblastoma cells. **A.** Scatter plots of γH2AX versus DNA content in A549 cells treated with radiation (6 Gy) and 10 μM 5-iodotubercidin (5-ITU), to validate screen results. Numbers indicate γH2AX median values. **B.** Bar charts showing γH2AX median values (relative to non-treated condition) and γH2AX_ratio_ (calculated as in Figure 5B) from multiple validation experiments in A549 cells (n=3). Error bars represent the SEM, and each dot represents an individual experiment. *p<0.05, **p<0.01, ***p<0.001 (two tailed Student t test). **C.** Similar as in B for U251 glioblastoma cells treated with radiation (6 Gy) and 5-iodotubercidin (5-ITU; 500 nM) for 30 min or 24 hours. The γH2AX_ratio_ was calculated as (γH2AX_6Gy,24h_ –γH2AX_0Gy,24h_)/ (γH2AX_6Gy,30min_ – γH2AX_0Gy,30min_) (n=3). **D.** Similar as in C for T98G glioblastoma cells. (n=3). **E.** Clonogenic survival of U251 (left) and T98G (right) cells treated with 5-iodotubercidin (500 nM) and the indicated doses of radiation. Error bars: SEM (n=3). *p<0.05, **p<0.01(two tailed Student t-test). **F.** Immunoblots of T98G (top) and A549 (bottom) treated with 5-iodotubercidin and radiation as indicated. All blots were run on the same gel. Uncropped blots are shown in Supplementary Figure S5C.

## Discussion

In this work we developed and optimized a large-scale flow cytometry screen to identify compounds that inhibit DNA repair after irradiation. The platform enables analysis of multiple endpoints using γH2AX as a marker for DSBs and phospho-H3 as a marker for mitosis. The ratio of γH2AX levels at 6 hours relative to 30 minutes after irradiation proved to be a robust parameter for detecting DNA repair inhibition, as a high ratio indicates persistent DNA damage at 6 hours. In addition, γH2AX levels at 30 minutes after irradiation served as an endpoint to identify compounds that interfere with radiation-induced phosphorylation of H2AX. By incorporating phospho-H3 staining, the screen also allowed detection of compounds that abrogate the radiation-induced G_2_ checkpoint. The screening approach successfully recovered known radiosensitizers and inhibitors of DNA repair while also identifying novel candidate compounds. Notably, 5-iodotubercidin radiosensitized glioblastoma cells despite previous reports describing radioprotective effects in normal brain tissue. These apparently divergent effects may reflect distinct mechanisms of action of 5-iodotubercidin in tumor versus non-proliferating normal cells, with radiosensitization in cancer cells potentially mediated by impaired DNA repair (this study), and radioprotection in normal brain tissue linked to modulation of adenosine metabolism and astrogliosis [25,26]. Altogether, these findings establish a robust flow cytometry-based screening platform for identifying DNA repair inhibitors and highlight 5-iodotubercidin and omipalisib as preclinical candidates for radiosensitizer development.

## DATA AVAILABLILITY STATEMENT

The data that support the findings of this study are available from the corresponding author upon reasonable request. The raw screening data are provided in Supplementary Table S2 (Excel file).

## Author contributions

Conceptualization: RGS, TS; Investigation: CN, GER, IAB, AEM, SH, LHS, LL, AGG, KK, VS; Investigation, supporting experiments: DS; Methodology: CN, TS, RGS; Project administration RGS: Supervision: RGS, GER, SH; Validation: CN, GER, VS, SH; Visualization: CN, LHS, AGG; Writing - original draft: RGS, CN; Writing - review & editing: all authors, Funding acquisition: RGS, DS.

## Additional information

The authors declare no competing interests.

