## Supplementary material for "A flow cytometry-based screening platform for identifying candidate radiosensitizers targeting DNA repair"

### Supplementary methods:

*Detailed description of flow cytometry screening procedure*

Compound libraries, including the Enzo Target and Pathway libraries (WNT, epigenetics, autophagy, and phosphatase inhibitors), the Selleck Chem Cambridge Cancer compound library, and the Biomol kinase inhibitor library, were obtained from the Chemical Biology Platform. Compounds were distributed in 384-well V-bottom plates (PerkinElmer, #6008590). Cell seeding, fixation, and staining were facilitated using a Precision XS microplate sample processor and an ELx405 Select microplate washer (BioTek).

A549 lung cancer or Reh leukemia cells were seeded at $10^5$ cells/100µl medium per well. The final compound concentration was 10µM, except one experiment with 1µM of the Selleck Chem Cambridge Cancer compound library. Three parallel plates were processed and analyzed together: all three containing compounds and two of them were treated with 6Gy after seeding. This dose produced a strong induction of γH2AX at 30 minutes followed by a marked reduction at 6 hours, providing an optimal window to detect inhibition of DNA repair. One of the irradiated plates was incubated for 30 minutes and the other two plates were incubated for 6 hours at 37ºC/5%$CO_2$ and thereafter pelleted by centrifugation. The cell pellets were fixed with 80µl 100% methanol per well. The plates were then stored at -20ºC until further analysis.

Before antibody staining, the plates were barcoded with Pacific Blue (PB) (Molecular Probes, P10163). Fixed plates were first pelleted by centrifugation. The 0Gy plate was then stained with the highest concentration of PB (0,2496 ng/µl), the 6 Gy 30 min plate by a lower concentration of PB (0,01248 ng/µl), and the 6 Gy 6 h plate was left without PB staining, in a 50µl volume each per well. The PB stained plates were incubated at room temperature in the dark for 20 minutes. PB staining was blocked by addition of 40µl of a PBS/1% FBS solution per well. Plates were pelleted by centrifugation, and an additional 90µl of PBS(1x)/FBS (1%) solution was added per well. Plates were pelleted by centrifugation. Further, 20µl (per well) of PBS/1% FBS solution was added to two of the three plates and the contents of these two plates were transferred into the corresponding wells of the third plate. The resulting plate was pelleted by centrifugation.

For the DNA damage repair endpoint, cells were stained with an antibody to the DNA damage marker γH2AX. For simultaneous assessment of $G_2$ checkpoint abrogation (in Reh cells), cells were co-stained with an antibody to the mitotic marker phospho-H3. Cells were incubated for 1 hour at room temperature in the dark with 25µl (per well) of anti-phospho-H2A.X (Ser139) clone JBW301, FITC conjugated (mouse monoclonal) antibody (16-2020A, Millipore) and anti-phospho-H3 (Ser10), PE-conjugated (Rabbit monoclonal) antibody (#5764, BioNordika), both diluted 1:1000 in PBS containing 0.2% Tween-20 (Sigma) and 4% milk powder. Cells were washed with 65µl of a PBS(1x)/FBS(1%) solution per well. The cell pellet was resuspended in 90µl (per well) of the DNA stain FxCycle FarRed solution (PBS, RNAse A). The stained plates were stored at 4°C in the dark overnight.

Flow cytometry analysis was performed with an LSR II flow cytometer (BD Biosciences) equipped with a BD High Throughput Sampler using the FACS Diva Software version 6.1.3 (BD Biosciences) during acquisition. The FlowJo software (FlowJo, LLC) was used during analysis. The three populations of the different treatments were separated and defined based on the Pacific Blue signal. The median γH2AX signal was obtained for each sample, as well as the mitotic population with strong phospho-H3 in a defined area above the $G_2$ population.

To identify possible candidate hits regarding the γH2AX endpoint in the screen we calculated the ratio of γH2AX level at 6 hours relative to the γH2AX level at 30 minutes for all compounds ($\gamma H2AX_{ratio} = (\gamma H2AX_{6Gy,6h} - \gamma H2AX_{0Gy,6h}) / (\gamma H2AX_{6Gy,30min} - \gamma H2AX_{0Gy,6h})$). The subtraction of values for non-irradiated samples ($\gamma H2AX_{0Gy,6h}$) was included to deduct possibly increases in γH2AX caused by the compounds alone, in the absence of irradiation. To define candidate hits with clearly higher $H2AX_{ratio}$ than the majority of samples, a threshold was set by eye in $H2AX_{ratio}$ histograms (see e.g. Figure 2B, stippled lines). For the $G_2$ abrogation endpoint we compared the percentage of phospho-H3 positive mitotic cells for the 6 Gy 6h sample to the percentage for the 6h 0 Gy sample, for all compounds. Control samples had about 2% mitotic cells with 0Gy and <0.05% mitotic cells with 6 Gy. Compounds showing more than ~ 0.5% mitotic cells at 6 hours after 6Gy, while not showing an increased percentage of mitotic cells in the corresponding non-irradiated sample (0 Gy, 6h), were identified as candidate hits (see e.g. Figure 3C).

*Immunofluorescence*

T98G cells seeded on 10mm glass coverslips (in 35 mm dishes) were treated with 1µM 5-iodotubercidin or DMSO (0.01%) control ~ 30 minutes prior to irradiation with 6 Gy. Cells were fixed in formalin 24 hours later. For immunofluorescence, the cells were permeabilized in PBS containing 0.5% Triton X-100 and incubated for 1 hour at room temperature with antibodies against γH2AX (Millipore, clone JBW301, 1:400) and phospho-53BP1 (Cell Signaling Technology, 2675S, 1:100) for 1h at room temperature, followed by Alexa Fluor 488- and 568-conjugated secondary antibodies (Invitrogen, 1:1000) for 30 minutes. DNA was stained with Hoechst 33342 (0.6µg/ml in PBS), and samples were mounted with ProLong Diamond Antifade

Mountant (Invitrogen, P36965). Images were acquired using a spinning-disk confocal microscope (Nikon ECLIPSE Ti2-E CrestOptics X-Light V3) with a 60x oil-immersion objective.Channel adjustments and image montages were created in Fiji (ImageJ) version 1.54f.

**Supplementary table 1:**

| **Antibody** | **Source** | **Identifier** | **Dilution used** | **Application** |
|---|---|---|---|---|
| anti-phospho-H2A.X (Ser139) clone JBW301, FITC conjugated | Millipore | 16-2020A | 1:1000 | Flow cytometry screening |
| anti-phospho-H3 (Ser10), PE-conjugated | BioNordika | #5764 | 1:1000 | Flow cytometry screening |
| anti-phospho-H2A.X (Ser139) clone JBW301 | Millipore | 05-636 | 1:400 | Flow cytometry, Immunofluorescence |
| phospho-53BP1 | Cell Signaling Technology | 2675S | 1:100 | Flow cytometry, Immunofluorescence |
| Alexa Fluor 488 Donkey Anti-Mouse IgG | Life Technologies | A21202 | 1:1000 | Flow cytometry, Immunofluorescence |
| Alexa Fluor 568 Donkey anti-Rabbit IgG | Life Technologies | A10042 | 1:1000 | Flow cytometry, Immunofluorescence |
| DNA-PK phospho s2056 | Abcam | ab18192 | 1:400 | Western blotting |
| Chk1 phospho-Ser345 | Cell Signaling | 2348 | 1:400 | Western blotting |
| Chk1 phospho-Ser317 | Cell Signaling | 2344 | 1:400 | Western blotting |
| Chk1 (DCS-310) | ThermoFisher | MA1-9108[illegible] | 1:200 | Western blotting |
| RPA phospho(S4/S8) | Nordic Biosite | A300-245 | 1:1000 | Western blotting |
| Phospho-ATM (Ser1981) (10H11.E12) | Cell Signaling | 4526 | 1:1000 | Western blotting |

### Figure legends, supplemental figures.

**Figure S1. Screening of 357 compounds from the Enzo Pathway and BioMol Kinome libraries in Reh cells using phospho-H3 as the endpoint.**

**A.** Representative scatter plots of phospho-H3 versus DNA content for cells treated with radiation, with and without the proteasome inhibitor MG-132. Numbers indicate the percentage of mitotic cells, defined as cells with strong phospho-H3 signal and $G_2$/M DNA content. Data are from the second screen in Figure 2B, which included co-staining for phospho-H3 (S10).

**B.** Histograms showing the percentage of mitotic cells for each compound (red) and radiation-only controls (blue). Top: non-irradiated samples (0 Gy, 6 h); bottom: irradiated samples (6 Gy, 6 h).

**C.** Scatter plot of mitotic cell percentages at 6 h in irradiated versus non-irradiated samples, corresponding to the data in (B). Compound-treated samples are shown in red and controls in blue. Candidate hits, defined as compounds that abrogate the radiation-induced $G_2$ checkpoint without increasing mitotic entry in non-irradiated cells, are highlighted (oval).

**D.** Top: list of candidate hit compounds identified in (C). Bottom: Bar chart showing mitotic cell percentages at 6 h for nine control samples (black/gray) and the four candidate hits.

**Figure S2. Comparison of screen results in Reh and A549 cell lines.**

Scatter plots showing the parameter $\gamma H2AX_{ratio}$ in Reh versus A549 cells, for the screens with 357 compounds from the Enzo pathway and Biomol kinome libraries (results in Figure 2C and Figure 3B). Quadrant Q1 contains candidate hits scoring in both cell lines, Q2 and Q4 contain candidate hits scoring in one of the cell lines only, and Q3 contains all the control samples and compounds that did not score in any of the cell lines. Top panel: The results of the 1st screen in Reh cells plotted against the results of the 1st or 2nd screen in A549 cells, respectively. Bottom panel: The results of the 2nd screen in Reh cells plotted against the results of the 1st or 2nd screen in A549 cells, respectively.

**Figure S3. Analysis of radiation-induced H2AX (30 min after treatment)**

**A.** Histograms showing γH2AX level at 30 min after irradiation (6 Gy), for all compound samples (red) and controls without compound (blue). Results are from the same screens as in Figure 4.

**B.** Comparison of results for the two screens in A. The γH2AX level at 30 min after 6 Gy in the $1^{st}$ versus the $2^{nd}$ screen is shown for each compound (red) and control sample (blue). Quadrant Q3 contains compounds that gave lower induction of γH2AX at 30 min after 6 Gy in both screens. Quadrants Q2 and Q4 contain compounds that gave lower induction of γH2AX in one of the screens only, and Q1 contains all the control samples and the compounds that did not give lower induction of γH2AX after 6 Gy in any of the screens.

**C.** List of compounds giving lower induction of γH2AX at 30 min after 6 Gy in both screens (compounds in quadrant Q3 in B).

**Figure S4 Analysis of omipalisib- or 5-iodotubercidin-treated cells**

**A.** DNA histograms showing cell cycle profiles for the experiment in figure 5A.

**B.** Similar as in A for the experiment in figure 6A (5-ITU, 5-iodotubercidin).

**C.** Quantification of phospho-53BP1 values from the same experiments as in figure 6D (T98G cells). The phospho-$53BP1_{ratio}$ was calculated similarly as for the $\gamma H2AX_{ratio}$ (n=3). The error bars represent the SEM, and each dot represents an individual experiment. *$p<0.05$, **$p<0.01$, ***$p<0.001$ (two tailed Student t test).

**D.** Representative immunofluorescence images of phospho53BP1 and γH2AX in T98G cells at 24h after treatment.

**Figure S5 Uncropped Western blots**

Uncropped blots corresponding to the results shown in Figure 5E (**A**), Figure 5F (**B**), and Figure 6F (**C**) are presented. The regions used for the cropped images in the main figures are indicated in red. Membranes were cut into defined regions prior to antibody incubation, and each membrane was probed with a specific antibody. For some blots, stripping and re-probing were performed, as indicated in the figure. For one older Western blot experiment (B), the uncropped image contains all lanes of the original gel; however, molecular weight markers are not present in the archived file. Band identity was therefore determined based on the known position of the membrane cut, together with antibody specificity.

Figure S1

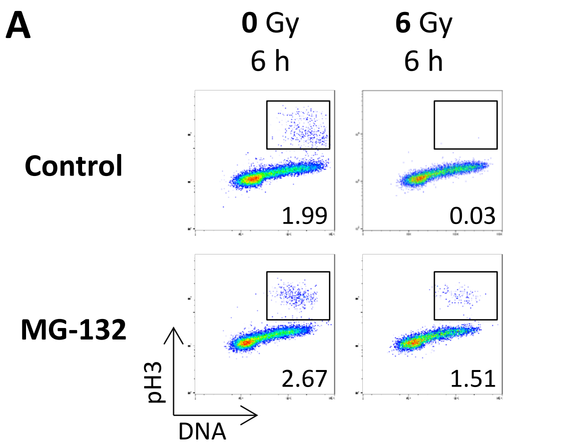


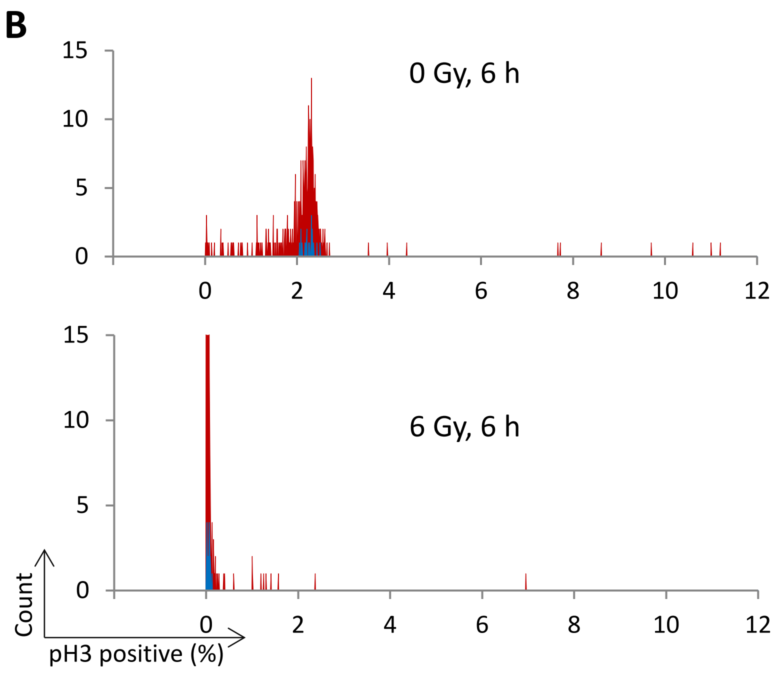


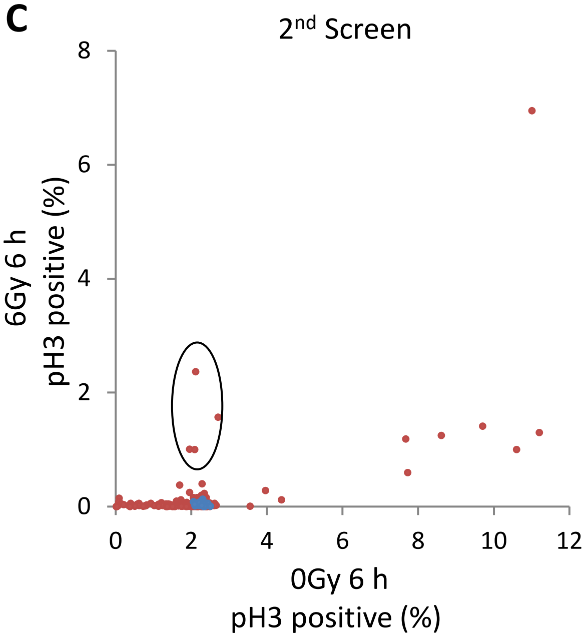


D

| Compound name | Target/Function | pH3 positive (%) 0Gy 6h | pH3 positive (%) 6Gy 6h |
|---|---|---|---|
| Sanguinarine chloride | TNF, STAT3, MMP-2, MMP-9 | 2,11 | 2,37 |
| MG-132 | Proteasome | 2,70 | 1,57 |
| Pimozide | Ca2+ channel | 1,95 | 1,01 |
| Rapamycin | mTOR | 2,08 | 1,00 |
| All controls | | 2,25 | 0,05 |

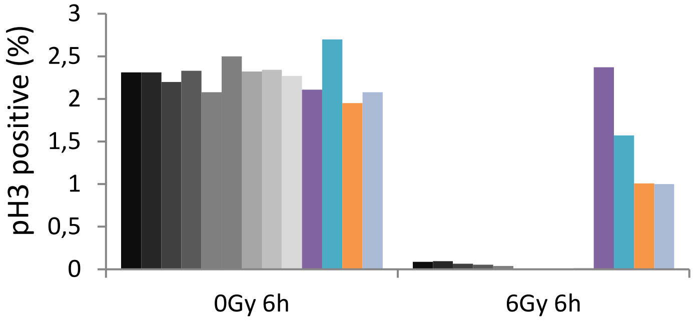

Figure S2

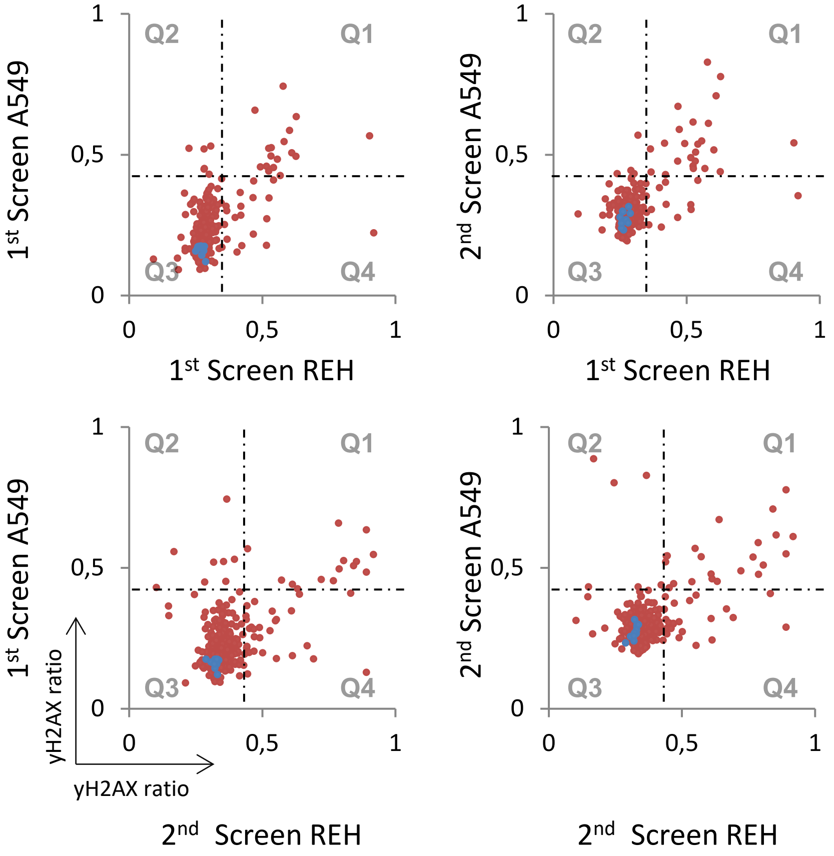
Q1
Q2
Q3
Q4
1st Screen A549
2nd Screen A549
1st Screen REH
2nd Screen REH
yH2AX ratio
yH2AX ratio
0
0,5
1

Figure S3

**A**

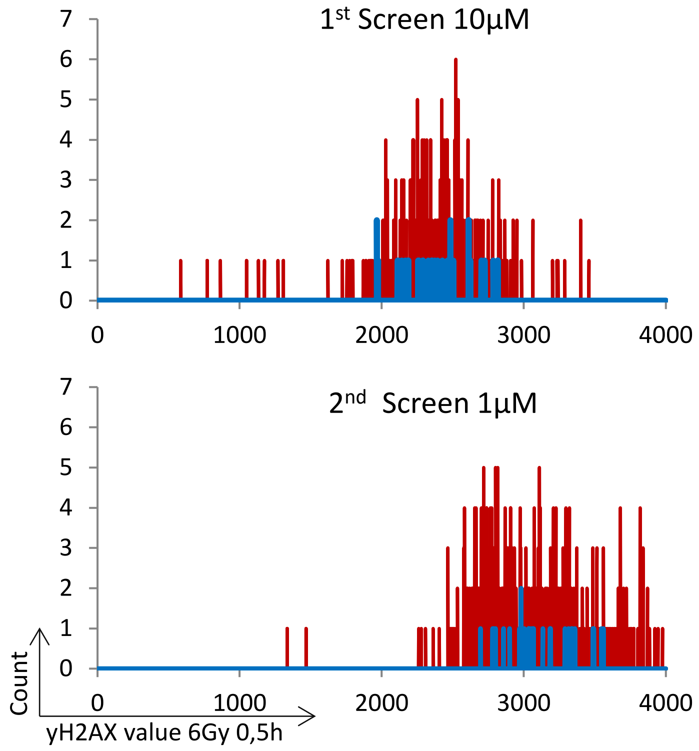


**B**

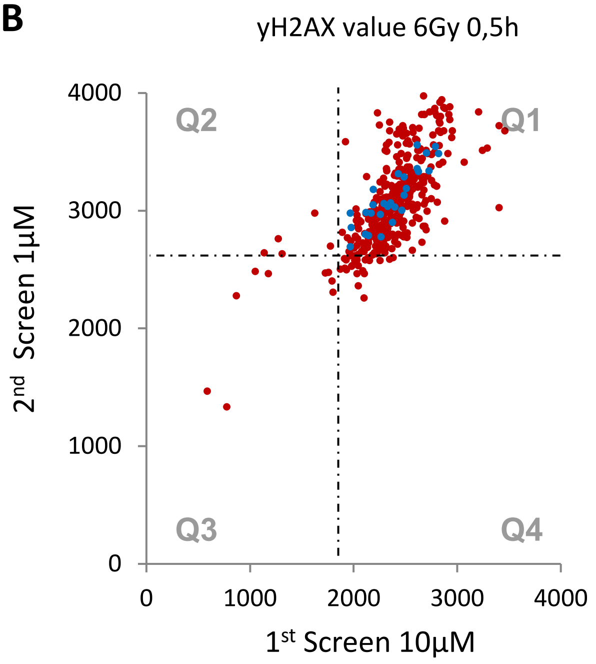


**C**

| Compound name | Target/Function | yH2AX value 6Gy 0,5h | |
|---|---|---|---|
| | | 10µM Screen | 1µM Screen |
| KU-60019 | ATM | 772 | 1335 |
| CP-466722 | ATM | 1174 | 2466 |
| PI-103 | DNA-PK, PI3K, mTOR | 1723 | 2471 |
| Torin 2 | mTOR | 586 | 1468 |
| INK 128 | mTOR | 1049 | 2485 |
| YM201636 | PI3K | 1756 | 2476 |
| GSK2126458 | PI3K, mTOR | 865 | 2278 |
| PIK-93 | PI3K, VEGFR | 1797 | 2309 |
| Phloretin (Dihydronaringenin) | PKC | 1789 | 2405 |
| Average of Controls | | 2368 | 3116 |

Figure S4

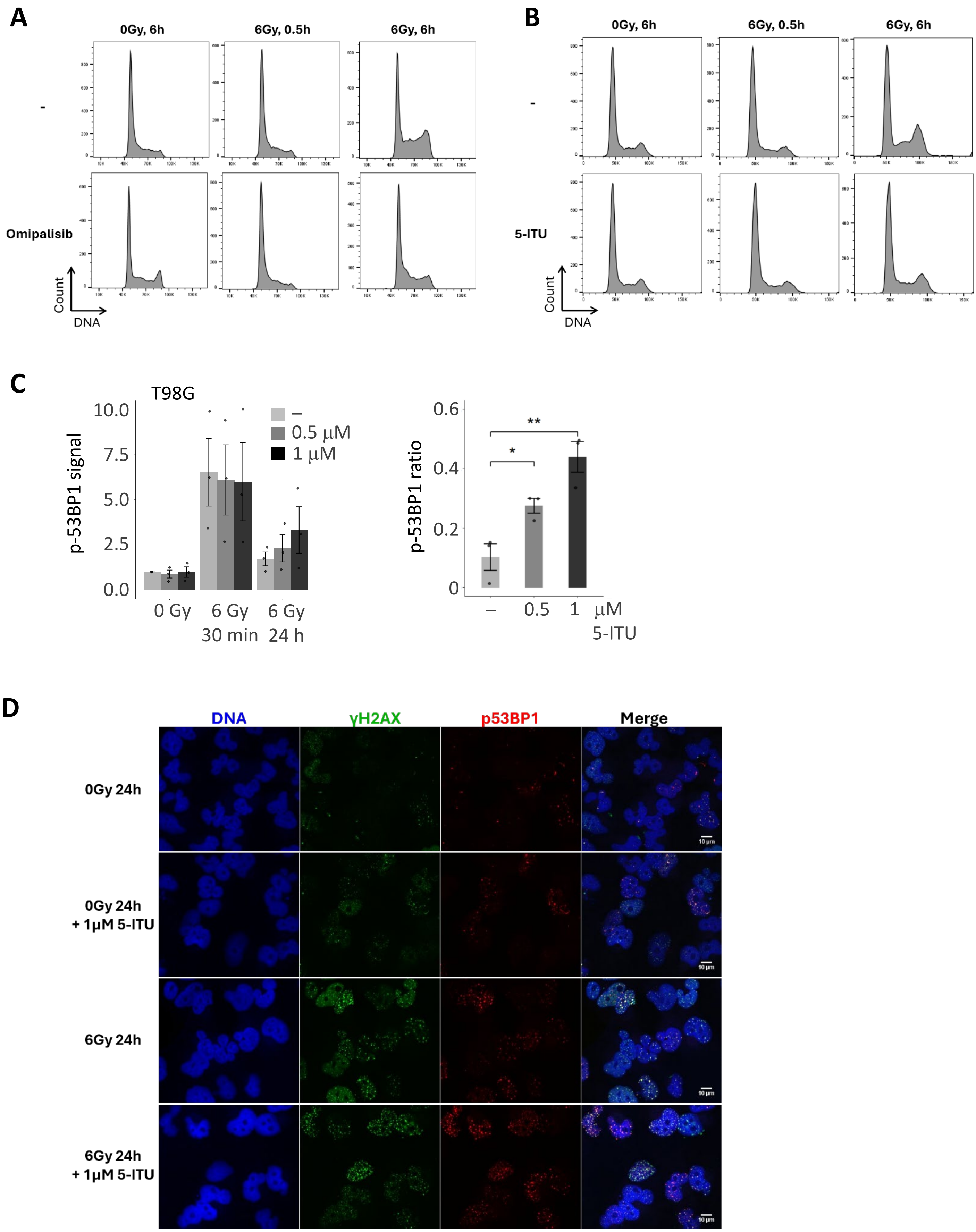

A
0Gy, 6h
6Gy, 0.5h
6Gy, 6h
-
Omipalisib
Count
DNA
B
0Gy, 6h
6Gy, 0.5h
6Gy, 6h
-
5-ITU
Count
DNA
C
T98G
p-53BP1 signal
10.0
7.5
5.0
2.5
0.0
–
0.5 μM
1 μM
0 Gy
6 Gy 30 min
6 Gy 24 h
p-53BP1 ratio
0.6
0.4
0.2
0
**
*
–
0.5
1
μM 5-ITU
D
DNA
γH2AX
p53BP1
Merge
0Gy 24h
0Gy 24h + 1μM 5-ITU
6Gy 24h
6Gy 24h + 1μM 5-ITU
10 μm

Figure S5, Uncropped Western blots

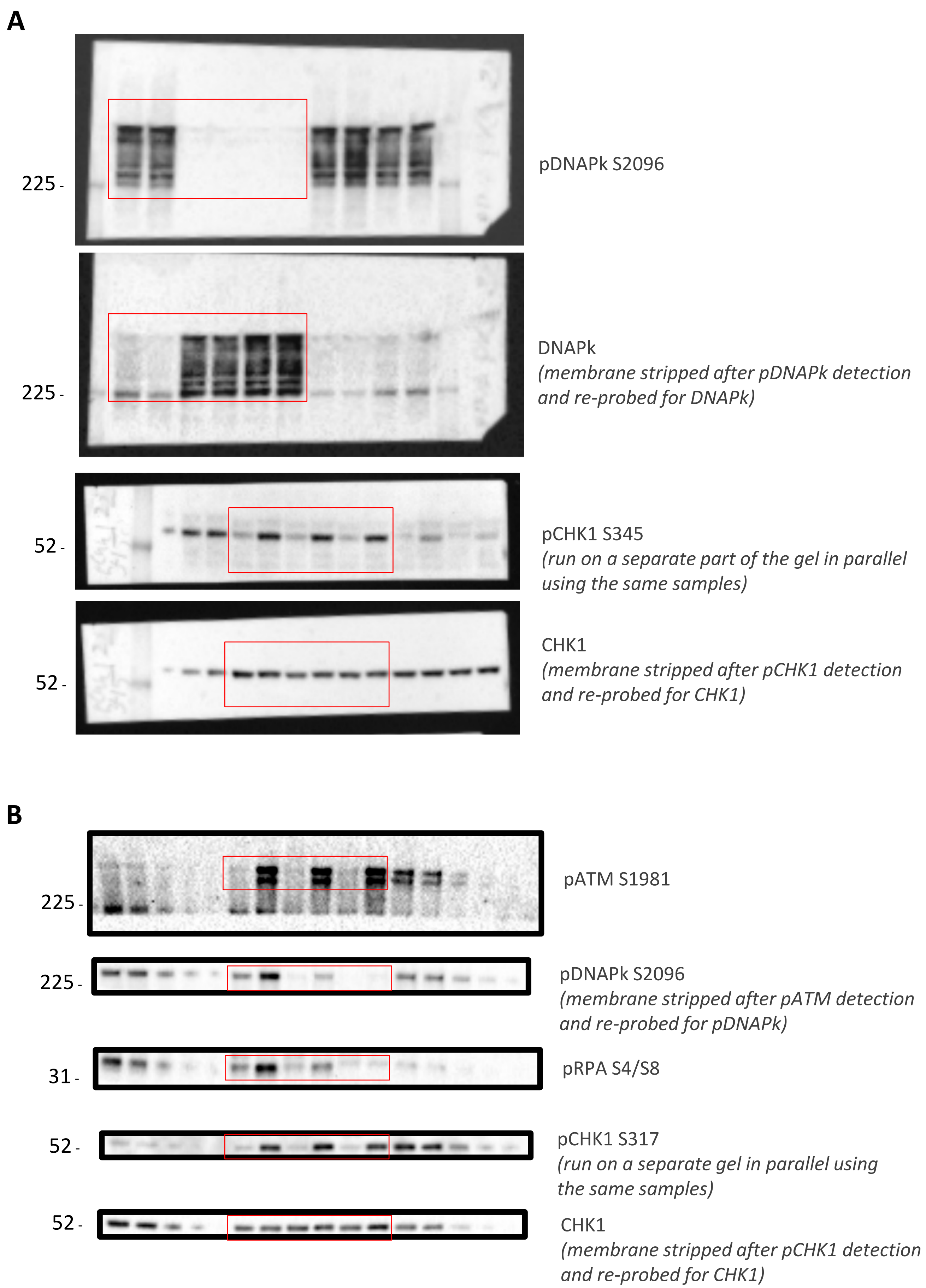

Figure S5, Uncropped Western blots

**C**

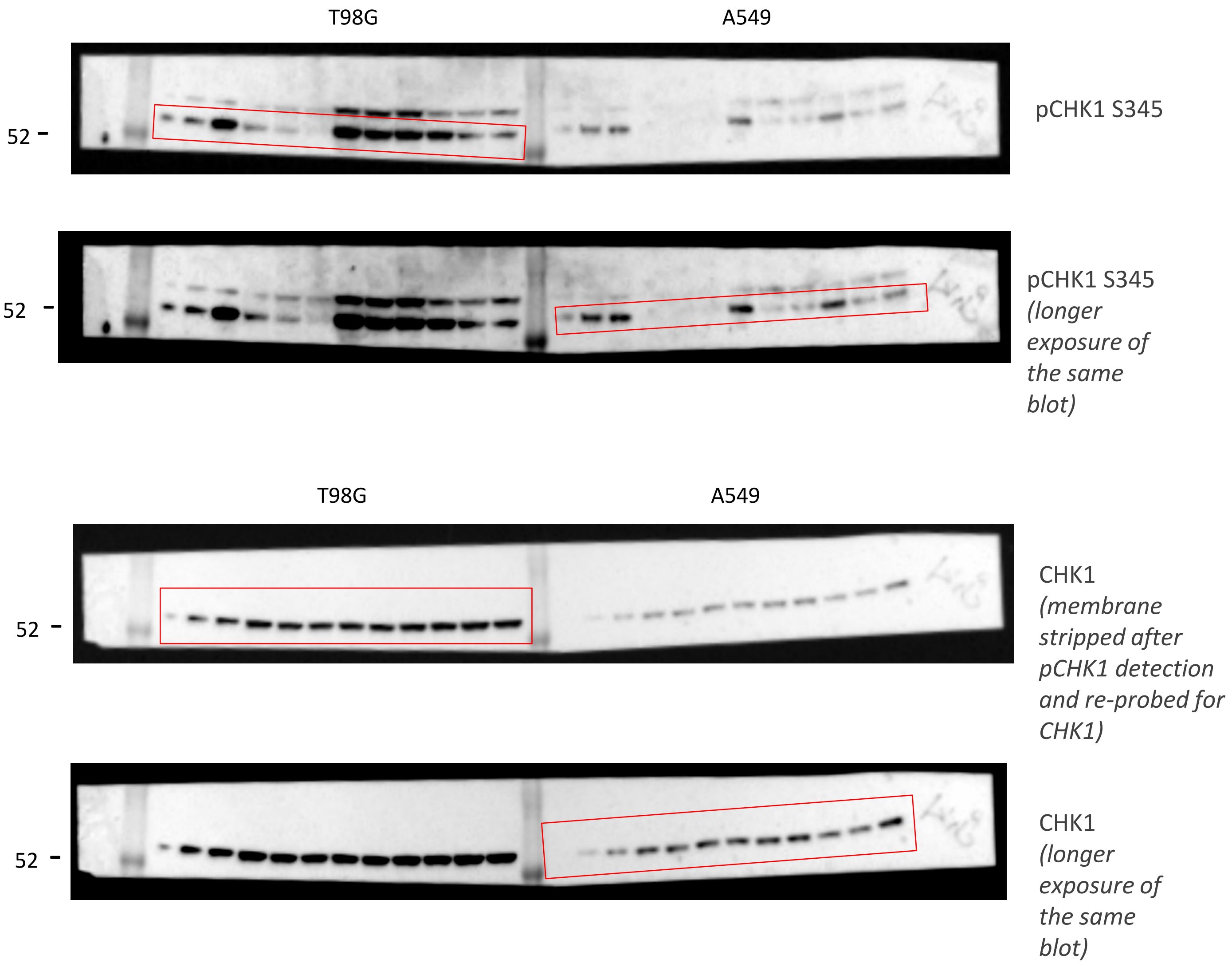
